# Prediction of plant resistance proteins using alignment-based and alignment-free approaches

**DOI:** 10.1101/2024.07.22.604583

**Authors:** Pushpendra Singh Gahlot, Shubham Choudhury, Nisha Bajiya, Nishant Kumar, Gajendra P. S. Raghava

**Affiliations:** Department of Computational Biology, Indraprastha Institute of Information Technology, Okhla Phase 3, New Delhi-110020, India

**Keywords:** PDR, Alignment-based approach, Machine learning, Ensemble method

## Abstract

Plant Disease Resistance (PDR) proteins are critical in identifying and killing plant pathogens. Predicting PDR protein is essential for understanding plant-pathogen interactions and developing strategies for crop protection. This study proposes a hybrid model for predicting and designing PDR proteins against plant-invading pathogens. Initially, we tried alignment-based approaches, such as BLAST for similarity search and MERCI for motif search. These alignment-based approaches exhibit very poor coverage or sensitivity. To overcome these limitations, we developed alignment-free or machine learning-based methods using compositional features of proteins. Our machine learning-based model, developed using compositional features of proteins, achieved a maximum performance AUROC of 0.92. The performance of our model improved significantly from AUROC of 0.92 to 0.95 when we used evolutionary information instead of protein sequence. Finally, we developed a hybrid or ensemble model that combined our best machine learning model with BLAST and obtained the highest AUROC of 0.98 on the validation dataset. We trained and tested our models on a training dataset and evaluated them on a validation dataset. None of the proteins in our validation dataset are more than 40% similar to proteins in the training dataset. One of the objectives of this study is to facilitate the scientific community working in plant biology. Thus, we developed an online platform for predicting and designing plant resistance proteins, “PlantDRPpred” (https://webs.iiitd.edu.in/raghava/plantdrppred).

**Highlights:**

- Development of a Machine-learning model for resistance protein prediction.
- Used alignment-based and alignment-free ensemble methods.
- Web server development and standalone package.
- Prediction and design of PDR proteins.

## 1. Introduction

A wide range of pathogens, including fungi, bacteria, nematodes, viruses, protozoa, and insects, can destroy or affect the growth of plants. Over the years, plants have evolved complex and dynamic immune systems essential for survival, growth, and development. Broadly, the plant immune system can be categorized into two main types: cell surface or pattern-triggered immunity (PTI) and intracellular or effector-trigger immunity (ETI). Pattern recognition receptors (PRRs) are specialized proteins that play a pivotal role in PTI by recognizing and responding to pathogen-associated molecular patterns (PAMPs) and damage-associated molecular patterns (DAMPs) [1]. Disease resistance proteins, products of Resistance (R) genes, enable plants to recognize specific pathogen effectors in ETI. These effectors are molecules produced by pathogens to promote their growth within host tissues. The interaction between R genes and pathogen avirulence (Avr) gene products determines whether a plant is resistant or susceptible to a pathogen attack [2].

R genes in plants exhibit diverse structural features, reflecting the complex nature of plant-pathogen interactions. At the molecular level, R genes typically encode PDR proteins with conserved domains such as nucleotide-binding sites (NBS) and leucine-rich repeats (LRR). The NBS domain, which can be either N-terminal or centrally located within the protein, is involved in ATP or GTP binding and hydrolysis, essential for signal transduction in plant defense responses. The LRR domain, often located at the C-terminus, is responsible for protein-protein interactions and pathogen recognition, as it forms a versatile scaffold for binding to pathogen-derived molecules [3]. Additionally, many PDR proteins contain other domains, such as Coiled-Coil (CC) motifs or Toll/Interleukin-1 receptor (TIR), which mediate downstream signaling events upon pathogen recognition [4,5]. The modular structure of PDR proteins allows for diverse recognition specificities and signaling pathways, contributing to the robustness of plant immunity.

Current prediction methods such as NBSPred and NLR-parser utilize sequence similarity or domain-based approaches for predicting PDR proteins [6,7]. These methods may fail if new proteins are not similar to known annotated proteins. Several machine learning-based methods have been developed to overcome these challenges, like DRPPP, prPred, and stackRPred [8–10]. Most of these methods have been developed on old and outdated data. Thus, there is a need to develop highly accurate and reliable models for predicting PDR proteins. In this study, we have systematically attempted to create the largest possible dataset, PDR proteins from database PRGdb and non-PDR proteins from Swiss-Prot. To create a non-redundant dataset, we created clusters using CD-Hit at 40%; these clusters were partitioned into training and validation datasets. In contrast to existing methods, we developed a hybrid or ensemble method that combines alignment-based and alignment-free approaches (Figure 1).

**Figure 1:**
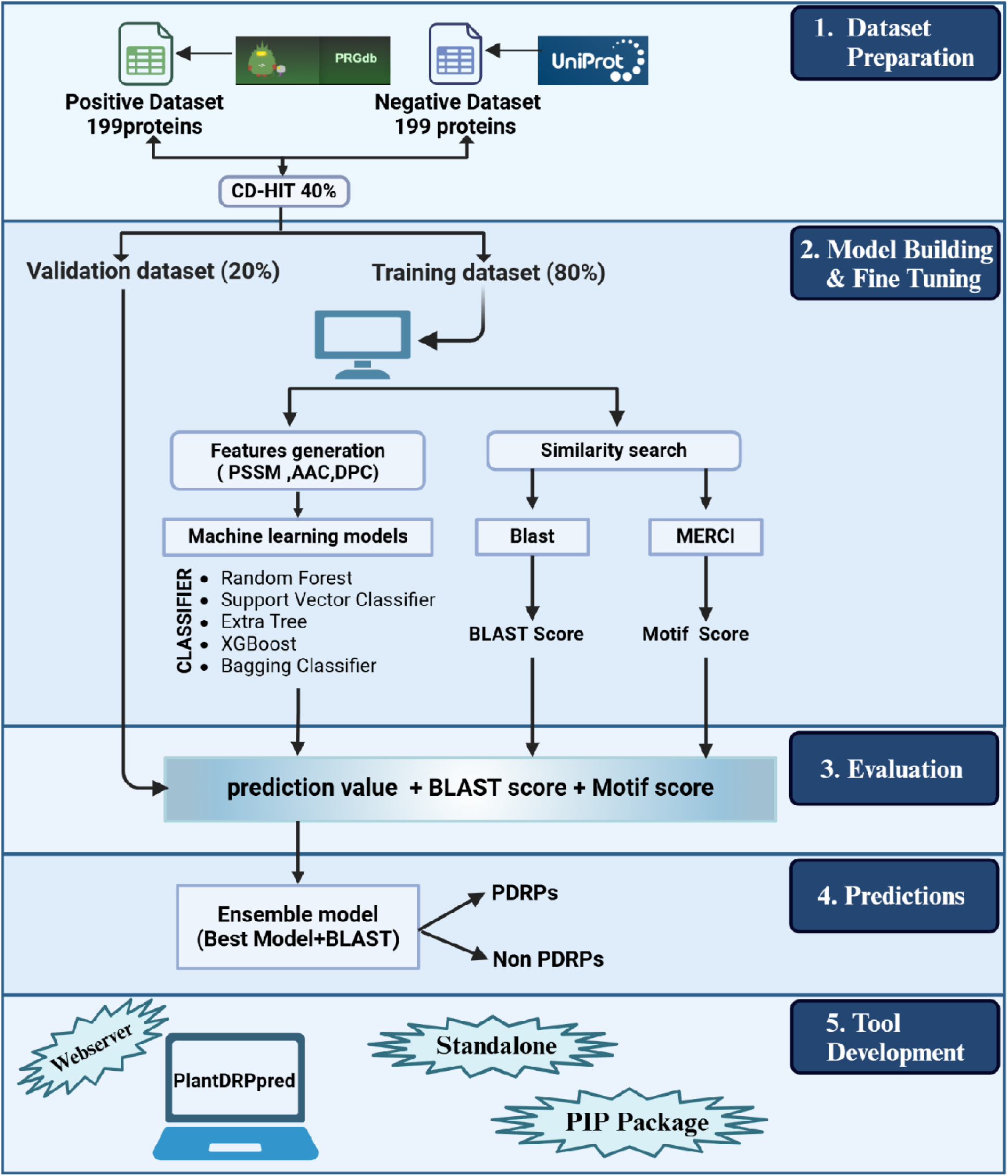
Workflow shows study architecture from data collection to web server development.

## 2. Materials and Methods

### 2.1. Dataset formation

In this study, we acquired 199 plant disease resistance (PDR) proteins from the PRGdb 4.0 database called positive proteins [11]. Similarly, we obtained non-PDR proteins from Swiss-Prot, called negative proteins [12]. Since the PDR protein sequences of Bryophyta and Angiosperm are available in positive sequences, we used sequences from the parent clade Embryophyta to generate the negative dataset, as it includes PDR proteins from both Bryophyta and Angiosperm. We specifically selected sequences that had been previously reviewed. Subsequently, we filtered this dataset to include sequences with lengths more than 63 and less than 1826. We randomly sampled 199 protein sequences from this filtered pool to constitute our negative dataset and ensure they are not involved in defense-related functions based on available information. To generate non-redundant subsets while preserving the number of sequences, we adopted an approach used in previous studies [13–16]. We used CD-HIT [17] software at a cut-off of 40% to create clusters for PDR proteins and non-PDR, where no two proteins have a sequence similarity of more than 40%; 118 clusters for PDR proteins and 186 clusters for non-PDR were obtained. The steps involved in creating a non-redundant dataset are illustrated in Figure 2.

**Figure 2:**
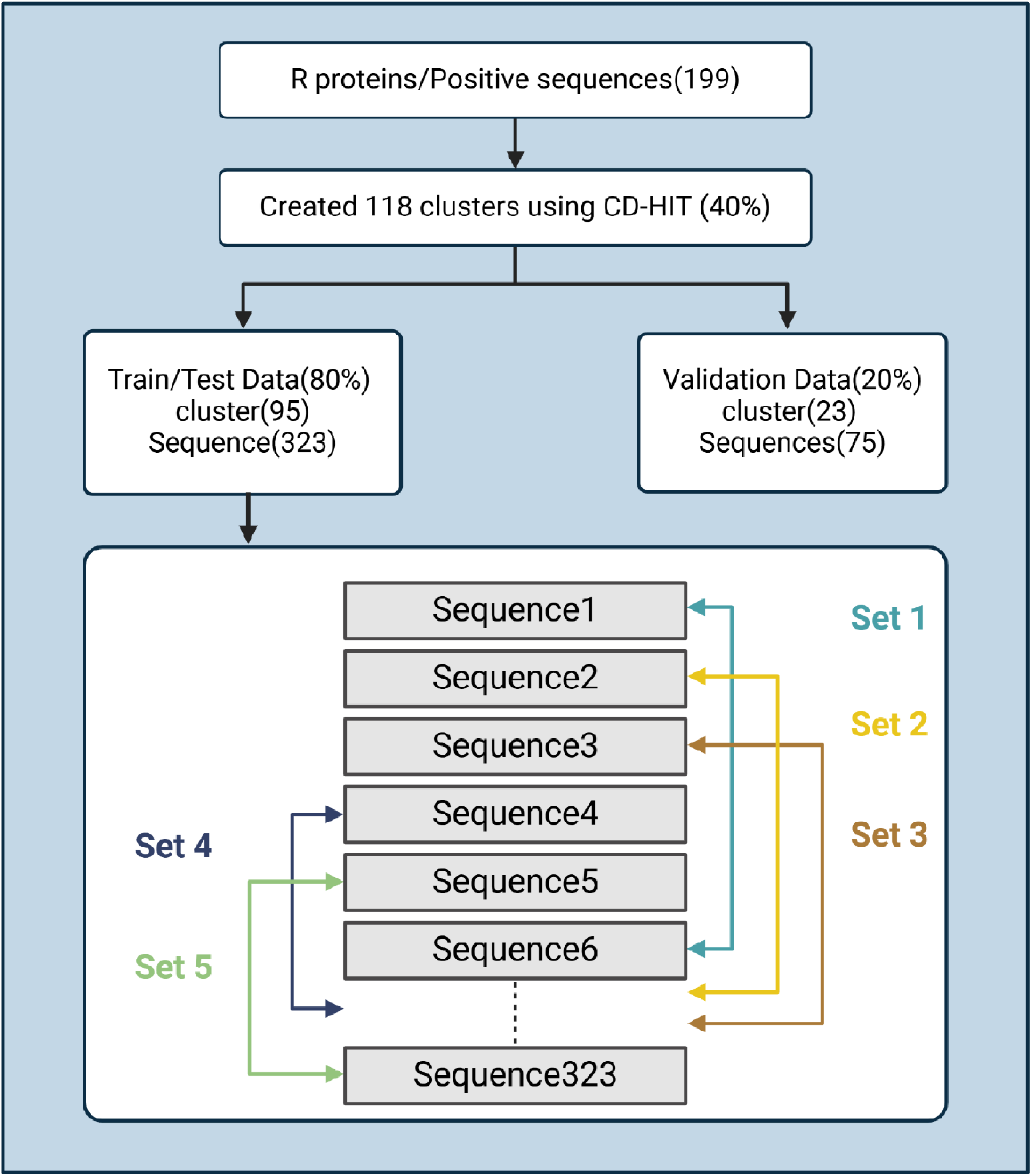
The flowchart illustrates the generation of non-redundant training, validation, and testing of datasets of PDR proteins. Initially, training and validation datasets are created by separating all clusters of PDR proteins. Subsequently, the training dataset’s sequences are divided into five subsets.

### 2.2. Feature generation

We utilized the tool Pfeature [18] to extract composition-based features and the POSSUM tool [19] to extract the Evolutionary feature (PSSM). Composition-based features include Amino Acid Composition (AAC) and Di-peptide Composition (DPC). AAC is represented as a 20-length vector, where each element signifies the fraction of a particular residue type within the sequence. In contrast, DPC is a 400-length vector that captures the occurrence of amino acid pairs within the protein sequence. The evolutionary features of proteins are recognized to provide vital insights beyond the primary sequence features [20]. These features are typically derived through the calculation of Position-Specific Scoring Matrix (PSSM) profiles using the Position-Specific Iterated Basic Local Alignment Search Tool (PSI-BLAST) [21]. The PSSM represents a matrix with dimensions of 20 × sequence length for protein or peptide sequences. However, to develop machine learning (ML) models, which require fixed-length vectors, we employed the composition of the PSSM profile as PSSM-400, a fixed-length vector containing 400 elements as evolutionary features [15].

### 2.3. Alignment based method

We employed BLAST (Basic Local Alignment Search Tool) [22] for similarity searches. First, we constructed a database using the training dataset. Then, we performed similarity searches against this database using sequences from the validation dataset. This approach allows us to compare sequences in the validation dataset against those in the training dataset, aiding in tasks such as classification or identification of similar sequences. Identifying functional motifs within protein sequences is crucial for functional annotation and distinguishing between positive and negative datasets. This study employed the Motif Emerging with Classes Identification (MERCI) program to identify motifs within PDR and non-PDR protein sequences [23]. We explored the extraction of motifs using various k-values. Afterward, we use parameter k=20 to extract motifs (Supplementary Table S2) that are exclusively present in PDR proteins and not non-PDR proteins.

### 2.4. Alignment Free methods

#### 2.4.1. Machine Learning Techniques

We implemented machine learning classifiers using the Python sci-kit learn package to build our prediction models. To enhance model performance, we fine-tuned hyperparameters on the training dataset with the help of sklearn’s GridSearch package. After determining the optimal hyperparameters, the best model was evaluated on the validation set. We used a five-fold cross-validation to execute the entire process and averaged the results across all five folds. Using optimized feature vectors, we developed prediction models with various classifiers, including Support Vector Machine (SVM), Extra Trees (ET), Random Forest (RF), Gradient Boosting (GB), and Bagging Classifier (BC).

### 2.5. Ensemble or Hybrid approach for classification

This study used a hybrid or an ensemble approach to augment the model’s predictive capabilities. This ensemble methodology adopts a weighted scoring mechanism, incorporating three distinct methods: (i) ML approach, (ii) similarity-based technique employing BLAST, and (iii) motif-based technique using MERCI. Scoring system in the BLAST approach, a weight of ‘+0.5’ was assigned for positive predictions, ‘-0.5’ for negative predictions, and ‘0’ for no hits using BLAST. A scoring system was devised for MERCI classification to assign values based on various conditions where, when a motif is found in a sequence, a score of ‘+0.1’ is assigned. Additionally, for each additional motif found in the same sequence, an extra ‘0.1’ is added. If a motif is found in a negative sequence, a score of ‘-0.1’ is added. If no motif is found in a sequence, a score of 0 is assigned.

### 2.6. Performance metrics calculation

The ML models employed in this investigation were assessed using a variety of performance metrics, encompassing parameters both dependent and independent of the threshold. The evaluation metrics include specificity, sensitivity, Matthew correlation coefficient (MCC), the area under the receiver operating characteristic curve (AUROC), and accuracy. Notably, AUROC is threshold-independent, while the remaining parameters, such as specificity, sensitivity, and MCC, are threshold-dependent and were optimized to identify the threshold yielding maximum values. These metrics have been widely employed in previous studies to gauge the performance of ML models [16,24,25]. The following equations 1–4 were used to calculate these:

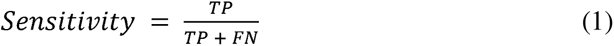

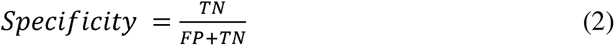

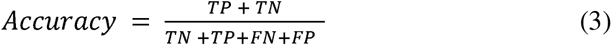

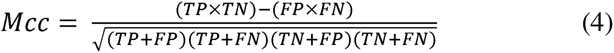

Where TP is a true positive, FP is a false positive, TN is a true negative, and FN is a false negative.

## 3. Results

### 3.1. Composition analysis

We calculated AAC for PDR and non-PDR proteins, as shown in Figure 3. We observed that amino acids leucine (L), serine (S), and isoleucine (I) are more abundant in PDR proteins. Similarly, amino acids alanine (A), proline (P), and glycine (G) are widespread in non-PDR proteins. We calculated AAC using the formula shown in Equation 5.

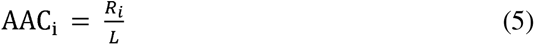

where is *AAC_i_* amino acid composition of residue type *i*; *R*_i_ is a number of amino acids of type *i*, and L is the length of the sequence.

**Figure 3:**
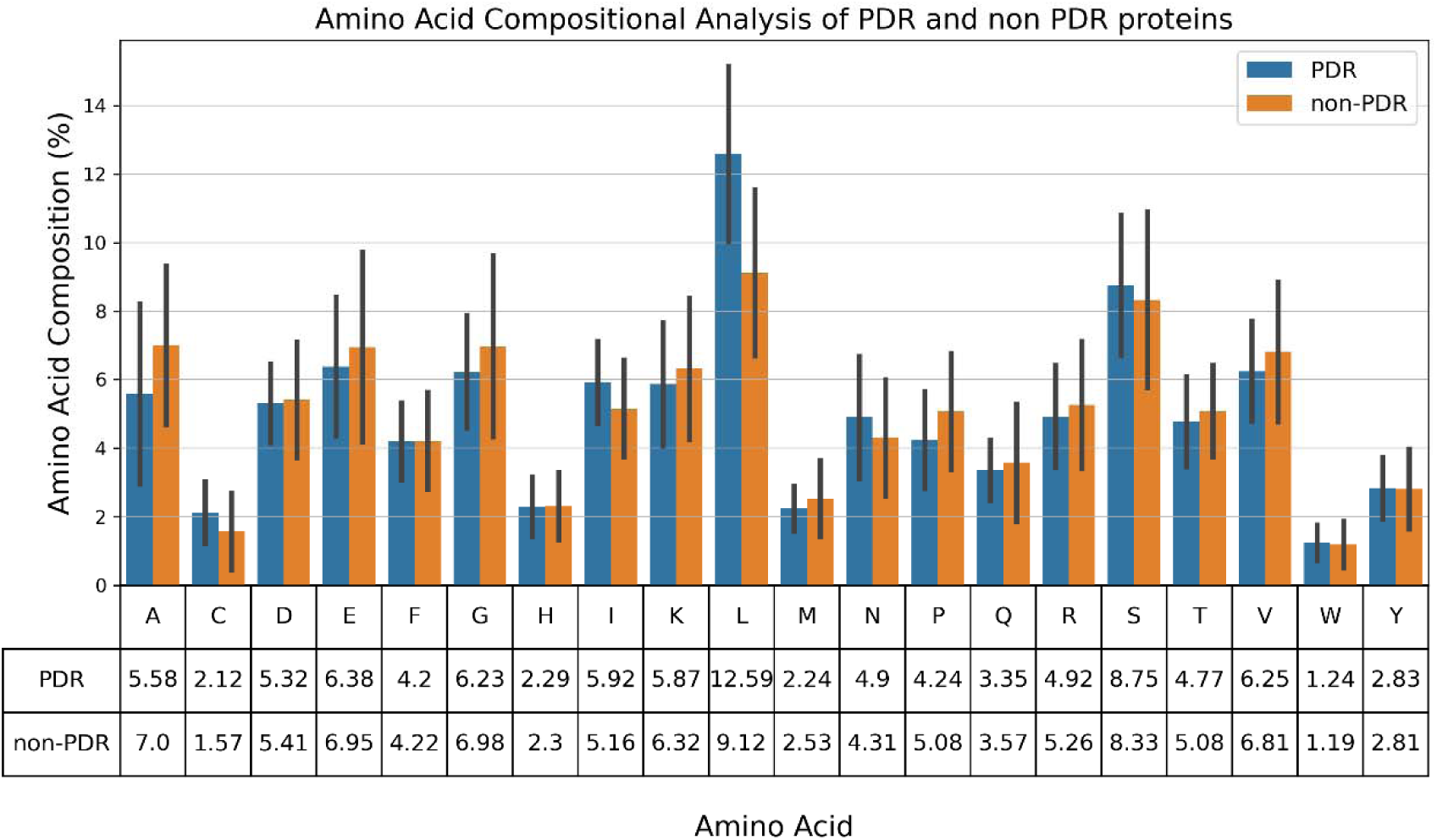
The percent amino acid compositional analysis for PDR or non-PDR proteins.

### 3.2. BLAST search

Initially, we utilized BLAST (version 2.15.0+) for similarity searches by constructing a database of training datasets. Subsequently, we employed this dataset to search for similarities within the validation sequences using various E-values. The performance of the BLAST module deteriorates when greater e-values are used, as BLAST permits random matches at higher e-values (see Table 1). Consequently, as the e-values increase, the number of hits also increases, but this negatively impacts performance because it allows for false hits.

**Table 1:** The hits obtained by employing BLAST on the validation datasets of PDR and non-PDR proteins at different e-values.

| Blast(e-value) | Correct Hits (PDRs) |  | Incorrect Hits (Non-PDRs) |  |
| --- | --- | --- | --- | --- |
|  | PDRPs | non-PDRPs | non- PDRPs | PDRPs |
| 1.00E-06 | 31 | 0 | 9 | 0 |
| 1.00E-05 | 31 | 0 | 9 | 0 |
| 0.0001 | 31 | 0 | 9 | 0 |
| 0.001 | 31 | 0 | 10 | 0 |
| 0.01 | 31 | 0 | 11 | 1 |
| 0.1 | 31 | 0 | 12 | 3 |
| 1 | 34 | 0 | 17 | 8 |
| 10 | 36 | 0 | 21 | 18 |
| 100 | 36 | 0 | 21 | 18 |

#PDRPs: Plant Disease Resistance proteins

### 3.3. Compositional-based Model

We utilized the AAC feature to construct ML models employing RF, ET SVC, XGB, and BC. The results are summarized in Table 2. The highest AUROC on the training set was achieved with SVC, reaching 0.92; similarly, SVC attained the highest AUROC of 0.91 on the validation dataset.

**Table 2:** The performance of different machine learning techniques-based models using the AAC feature of protein sequences.

| Models | Training Dataset |  |  |  |  | Validation Dataset |  |  |  |  |
| --- | --- | --- | --- | --- | --- | --- | --- | --- | --- | --- |
|  | Sen | Spec | Acc | AUROC | MCC | Sen | Spec | Acc | AUROC | MCC |
| **RF** | 0.81 | 0.86 | 0.83 | 0.91 | 0.68 | 0.86 | 0.77 | 0.81 | 0.88 | 0.63 |
| **SVC** | 0.81 | 0.87 | 0.84 | 0.92 | 0.68 | 0.75 | 0.87 | 0.81 | 0.91 | 0.63 |
| **ET** | 0.86 | 0.85 | 0.85 | 0.92 | 0.71 | 0.83 | 0.77 | 0.80 | 0.90 | 0.60 |
| **XGB** | 0.81 | 0.81 | 0.81 | 0.90 | 0.63 | 0.86 | 0.72 | 0.79 | 0.88 | 0.58 |
| **BC** | 0.79 | 0.85 | 0.82 | 0.91 | 0.64 | 0.86 | 0.79 | 0.83 | 0.88 | 0.66 |

Similarly, models were developed utilizing dipeptide composition along with various machine-learning techniques. Details of these models can be found in Supplementary Table S1. The best performance is the RF model with AUROC, which is 0.90 on the training set, and the validation dataset is 0.90.

### 3.4. PSSM feature-based Model

Previous studies have demonstrated the enhanced information provided by sequence profiles compared to individual sequences alone [16,26,27]. Therefore, in this study, we initially generated PSSM-400 composition profiles corresponding to each protein utilizing POSSUM software and used them as feature vectors to develop classification models. Similar to the AAC and DPC-based methods, we employed various classifiers, such as ET, RF, SVC, XGB, BC, etc. As depicted in Table 3, models based on evolutionary information exhibited a maximum AUROC of 0.96 on the training dataset. The PSSM profile was not generated for the sequence in the validation dataset. We used the model based on the second-best feature, AAC, to extract prediction values for that sequence. The maximum AUROC achieved was 0.96 on the validation set.

**Table 3:** The performance of different machine learning techniques-based models using the PSSM feature of protein sequences.

| Models | Training Dataset |  |  |  |  | Validation Dataset |  |  |  |  |
| --- | --- | --- | --- | --- | --- | --- | --- | --- | --- | --- |
|  | Sen | Spec | Acc | AUROC | MCC | Sen | Spec | Acc | AUROC | MCC |
| **RF** | 0.86 | 0.95 | 0.90 | 0.96 | 0.80 | 0.86 | 0.90 | 0.88 | 0.96 | 0.76 |
| **SVC** | 0.85 | 0.90 | 0.88 | 0.92 | 0.76 | 0.86 | 0.87 | 0.87 | 0.90 | 0.73 |
| **ET** | 0.86 | 0.96 | 0.91 | 0.95 | 0.82 | 0.86 | 0.95 | 0.91 | 0.95 | 0.82 |
| **XGB** | 0.88 | 0.90 | 0.89 | 0.94 | 0.78 | 0.86 | 0.87 | 0.87 | 0.95 | 0.73 |
| **BC** | 0.87 | 0.91 | 0.88 | 0.94 | 0.77 | 0.86 | 0.90 | 0.88 | 0.93 | 0.76 |

### 3.5. Ensemble Approach

The previous results indicate that both the similarity-based approach and machine learning-based models have their own advantages and disadvantages. Therefore, we endeavored to create a technique that integrates the strengths of both approaches. We explored hybrid methodologies, combining ML with BLAST scores and ML with BLAST and motif scores. we observed increased performance by combining the ML with the BLAST score using an E-value of 10^-3^. Specifically, when utilizing RF with features derived from PSSM, the AUROC improved to 0.98 on the validation dataset. Employing the ML model alongside exclusive positive motifs also resulted in an AUROC of 0.95 on the validation dataset. Combining machine learning with BLAST and motifs yielded an AUROC of 0.98 on the validation dataset. The performance of different ensemble approaches has been illustrated in Tables 4, 5, and 6.

**Table 4:**
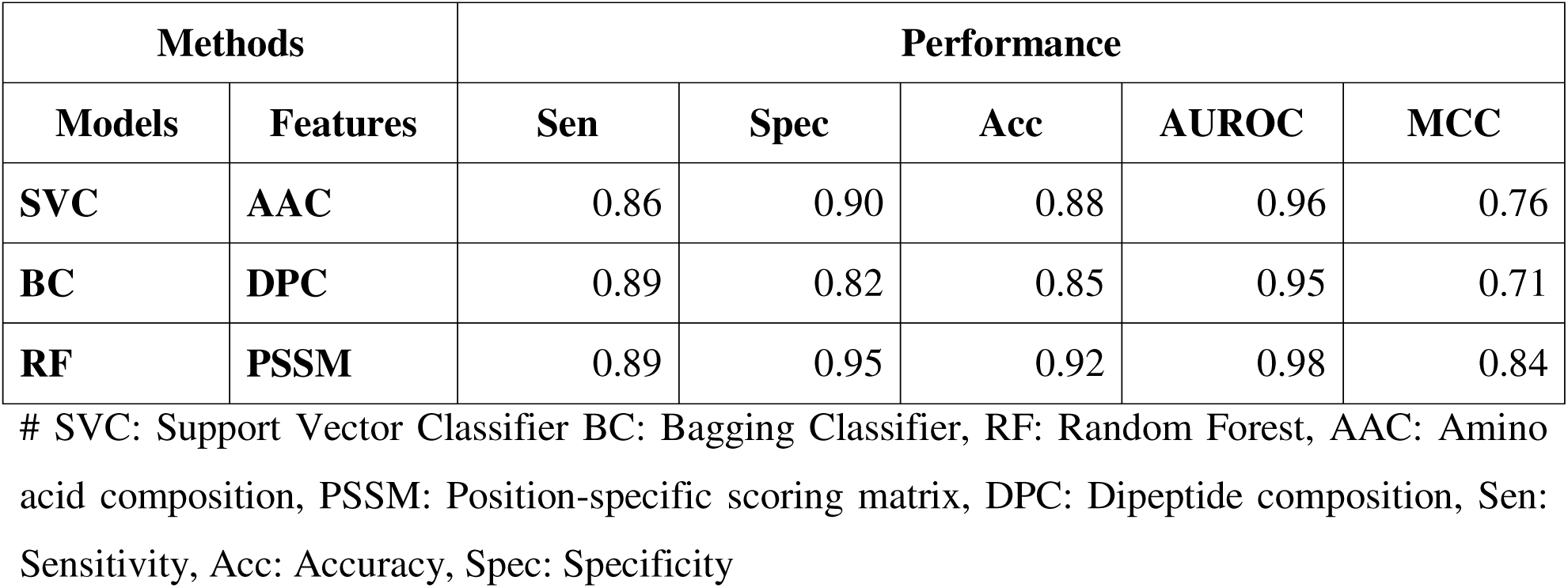
The performance of the hybrid method developed by combining machine learning and BLAST-based approach, evaluated on a validation dataset.

| Methods |  | Performance |  |  |  |  |
| --- | --- | --- | --- | --- | --- | --- |
| Models | Features | Sen | Spec | Acc | AUROC | MCC |
| **SVC** | **AAC** | 0.86 | 0.90 | 0.88 | 0.96 | 0.76 |
| **BC** | **DPC** | 0.89 | 0.82 | 0.85 | 0.95 | 0.71 |
| **RF** | **PSSM** | 0.89 | 0.95 | 0.92 | 0.98 | 0.84 |

**Table 5:**
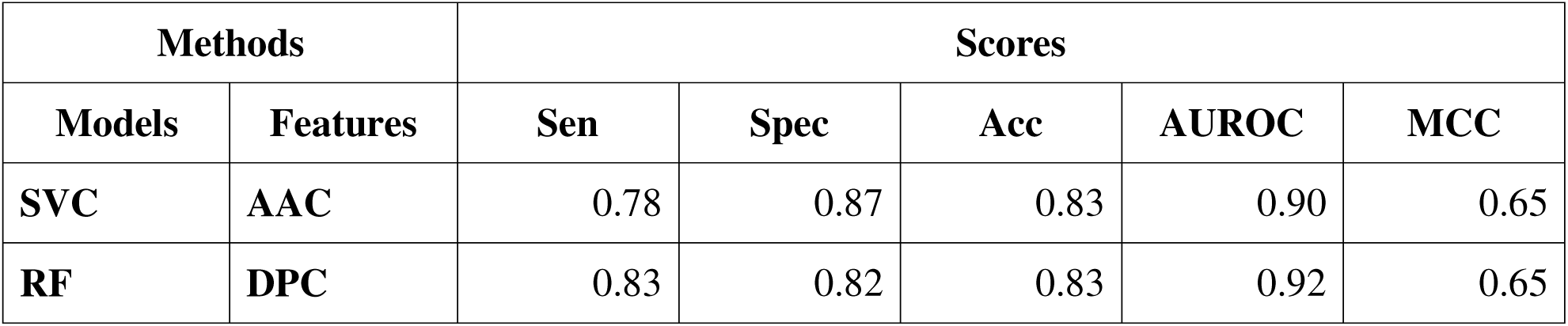
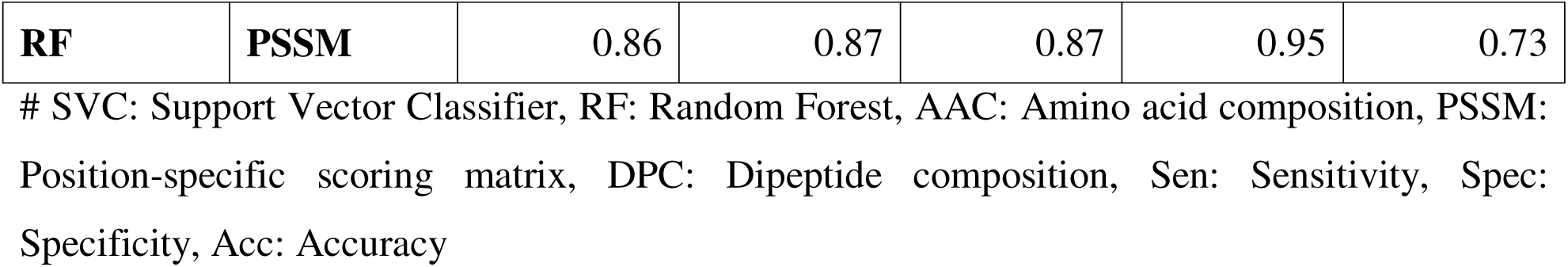
The performance of the hybrid method developed by combining machine learning and motif-based approach on a validation dataset.

**Table 6:** The performance of a hybrid method that combines machine learning, motif, and BLAST-based approaches on a validation dataset.

| Methods |  | Scores |  |  |  |  |
| --- | --- | --- | --- | --- | --- | --- |
| Models | Features | Sen | Spec | Acc | AUROC | MCC |
| **SVC** | **AAC** | 0.86 | 0.90 | 0.88 | 0.96 | 0.76 |
| **BC** | **DPC** | 0.89 | 0.82 | 0.85 | 0.95 | 0.71 |
| **RF** | **PSSM** | 0.89 | 0.95 | 0.92 | 0.98 | 0.84 |

## 4. Web Server Development for PDR Classification

We have constructed a web server called “PlantDRPpred” (https://webs.iiitd.edu.in/raghava/plantdrppred) for categorizing proteins into PDRPs (Plant Disease Resistance Proteins) or non-PDRPs groups, utilizing the top-performing model identified in our study. The server incorporates various modules: (i) Predict module, (ii) Design, (iii) Protein scan, and (iv) BLAST scan. Users can submit protein sequences for analysis, and the server will provide predictions and detailed reports generated by the incorporated modules.

## 5. Comparison with other tools

Our study evaluated the performance of various available tools for predicting R proteins. Some tools, NBSpred [6], DRPPP [8], and ResCap [28], StackRPred [10], are not available for evaluation. Available tools are trained on outdated datasets encompassing a limited range of plant disease-resistance protein classes, which restricts their ability to accurately predict all PDR protein classes. Our webserver, PlantDRPpred, overcomes these limitations and accurately predicts plant disease-resistance proteins. Table 7 represents the comparison of our model with the existing methods. The protein sequences in our validation dataset may already be present in the training dataset of existing tools. We have generated a dataset comprising proteins not utilized in the training or testing of existing methods and used that for benchmarking.

**Table 7:** Benchmarking of existing tools and tools proposed in this study on dataset not used in training of models.

| **Models** | **Sen** | **Spec** | **Acc** | **AUROC** | **MCC** |
| --- | --- | --- | --- | --- | --- |
| **prpred-DRLF** | 0.85 | 0.83 | 0.84 | 0.93 | 0.68 |
| **prpred** | 0.24 | 0.98 | 0.64 | 0.86 | 0.34 |
| **PlantDRPpred\*** | 0.89 | 0.95 | 0.92 | 0.98 | 0.84 |

## 6. Discussion

Plants encounter various types of biotic stress, including attacks from pathogens such as fungi, bacteria, insects, and viruses. These biotic stressors can severely affect plant health, reduce crop yields, and compromise food security. In response to these challenges, plants have evolved sophisticated defense mechanisms in which PDR plays a crucial role, recognizing specific pathogen-derived molecules and activating defense responses. This recognition triggers a cascade of signaling events to activate various defense mechanisms. Recent advancements in genomics and biotechnology have facilitated the identification and characterization of numerous PDR proteins across multiple plant species, but they are a time-consuming process. In the past few years, some approaches, such as DRPPP, prPred, and stackRPred, trained their model on older datasets having limited proteins and did not eliminate the redundancy of positive data. This study aims to establish an in-silico approach to predict plant disease resistance proteins using the ensemble model, so we retrieved 199 sequences from PRGdb4.0 as positive data. Negative data was extracted from UniProt, focusing on the Embryophyta clade since the positive data encompassed Angiosperm and Bryophyta. From the 5674 negative sequences retrieved from UniProt, we selected 199 sequences to generate a balanced dataset, ensuring that those involved in the defense system were excluded.

This study explored various methods to predict PDR proteins. We developed ML-based models to distinguish PDRs from non-PDRs by utilizing multiple features. These features included composition-based properties such as amino acid composition (AAC) and dipeptide (DPC), as well as evolutionary information-based properties derived from position-specific scoring matrices (PSSM). We employed a variety of classifiers, including SVC, RF, XGB, ET, and BC, to achieve this. Additionally, we used alignment-based approaches, such as BLAST and the Motif-search approach, to annotate protein sequences. However, these approaches demonstrated low sensitivity. Consequently, we adopted an ensemble approach that integrated the ML model with Blast with motif search, exploring all possible combinations of methods to improve the accuracy and reliability of our predictions. The highest performance was observed with a hybrid model combining the PSSM-based ML model and BLAST score. This hybrid approach was implemented in the freely accessible web server.

## 7. Conclusion

Our methodology for accurately predicting PDR proteins improves our capacity to recognize them better than the available methods. We used the ML model and BLAST approach for the final ensemble model. We have created a web server called PlantDRPpred and a standalone tool to assist researchers in identifying PDR proteins. If query sequences resembling PDR proteins are present, a prediction score based on similarity will be provided. We believe our work will contribute to the annotation of PDRs and provide valuable support for research in plant pathology.

## Data availability statement

The dataset used in this study is accessible from the “PlantDRPpred” web server at https://webs.iiitd.edu.in/raghava/plantdrppred/downloads.php. The source code can be obtained from https://github.com/raghavagps/PlantDRPpred.

## Author’s contribution

PSG and NK collected and processed the dataset. PSG developed computer programs and implemented the algorithms and prediction models. PSG and SC created the front-end and back-end of the Web server. PSG, NB, and GPSR wrote the manuscript. GPSR conceived and coordinated the project and provided overall supervision.

## Acknowledgments

The authors are thankful to the University Grants Commission (UGC), Council of Scientific and Industrial Research (CSIR), and Department of Science & Technology (DST) for their generous fellowships and financial support and to Indraprastha Institute of Information Technology Delhi for infrastructure. The authors would like to acknowledge the Department of Biotechnology (DBT) for the infrastructure grant awarded to the institute. Furthermore, they would like to acknowledge BioRender.com for creating the figures utilized in this work.

## Author’s Biography

1. Pushpendra Singh Gahlot is currently pursuing a Ph.D. in Computational Biology at the Department of Computational Biology, Indraprastha Institute of Information Technology, New Delhi, India.
2. Shubham Choudhury is currently pursuing a Ph.D. in Computational Biology at the Department of Computational Biology, Indraprastha Institute of Information Technology, New Delhi, India.
3. Nisha Bajiya is currently pursuing a Ph.D. in Computational Biology at the Department of Computational Biology, Indraprastha Institute of Information Technology, New Delhi, India.
4. Nishant Kumar is currently pursuing a Ph.D. in Computational Biology at the Department of Computational Biology, Indraprastha Institute of Information Technology, New Delhi, India.
5. Gajendra P.S. Raghava is currently working as a Professor and Head of the Department of Computational Biology, Indraprastha Institute of Information Technology, New Delhi, India.

## Abbreviation

PDR: Plant disease resistance
AAC: Amino acid composition
DPC: Dipeptide composition
PSSM: Position-specific scoring matrix
BLAST: Blast Local Alignment Search Tool
RF: Random forest
ET: Extra Trees
SVC: Support Vector Classifier
XGB: eXtreme Gradient Boosting
BC: Bagging Classifier
Sen: Sensitivity
Spec: Specificity
Acc: Accuracy
AUROC: Area Under the Receiver Operating characteristic Curve
MCC: Matthew Correlation Coefficient

