## Supplementary Table for "Prediction of plant resistance proteins using alignment-based and alignment-free approaches"

**Supplementary Table S1: The performance of different machine learning techniques-based models using the DPC feature of protein sequences.**

| **Models** | **Training Dataset** | | | | | **Validation Dataset** | | | | |
| --- | --- | --- | --- | --- | --- | --- | --- | --- | --- | --- |
|  | **Sen** | **Spec** | **Acc** | **AUROC** | **MCC** | **Sen** | **Spec** | **Acc** | **AUROC** | **MCC** |
| **RF** | 0.84 | 0.91 | 0.88 | 0.90 | 0.76 | 0.83 | 0.85 | 0.84 | 0.90 | 0.68 |
| **SVC** | 0.87 | 0.87 | 0.87 | 0.92 | 0.74 | 0.78 | 0.85 | 0.81 | 0.87 | 0.63 |
| **ET** | 0.83 | 0.92 | 0.88 | 0.90 | 0.76 | 0.86 | 0.85 | 0.85 | 0.88 | 0.71 |
| **XGB** | 0.79 | 0.82 | 0.80 | 0.88 | 0.61 | 0.83 | 0.74 | 0.79 | 0.86 | 0.58 |
| **BC** | 0.83 | 0.92 | 0.87 | 0.92 | 0.75 | 0.86 | 0.79 | 0.83 | 0.90 | 0.66 |

\# RF: Random Forest, ET: Extra Trees, SVC: Support Vector Classifier, XGB: eXtreme Gradient Boosting, BC: Bagging Classifier, Sen: Sensitivity, Spec: Specificity, Acc: Accuracy

**Supplementary Table S2: List of Exclusive Motif of PDR proteins using parameter K= 20 of MERCI.**

| **S. No.** | **Motifs** |
| --- | --- |
| 1 | G K T T L |
| 2 | G L P L |
| 3 | G K T T L A |
| 4 | K T T L A |
| 5 | L R K L |
| 6 | H L R Y |
| 7 | V L D D V |
| 8 | D D V W |
| 9 | H L R Y L |
| 10 | L D D V W |
| 11 | L L Y L |
| 12 | G L G K |
| 13 | G L P L A |
| 14 | L G K T |
| 15 | G L G K T |
| 16 | L I V L |
| 17 | V L D D V W |
| 18 | E G F V |
| 19 | G S R I |
| 20 | G I G K T |
| 21 | T S L E |
